# Transcranial magnetic stimulation to frontal cortex, unlike occipital stimulation, does not disrupt exogenous attention

**DOI:** 10.64898/2026.04.10.717799

**Authors:** Qingyuan (Rachel) Chen, Hsing-Hao Lee, Nina M. Hanning, Marisa Carrasco

## Abstract

Orienting covert attention to a target location improves performance across a wide array of visual tasks [1, 2]. Whereas fMRI studies have identified partially overlapping frontoparietal networks underlying endogenous (voluntary) and exogenous (involuntary) covert attention, these correlational methods cannot establish whether a given region is functionally necessary. Prior neurostimulation studies have established that early visual cortex (V1/V2) is critical for exogenous [3, 4] but not endogenous attention [5], whereas the right frontal eye field (rFEF+) is critical for endogenous attention [5]. Here, we used a combined psychophysical-TMS protocol to investigate whether rFEF+ is also required for exogenous attention. Participants performed an orientation discrimination task, in which a peripheral cue (valid, neutral, or invalid) preceded a target and a distractor stimulus. We applied two successive TMS pulses to rFEF+ during stimulus presentation and measured contrast-response functions (CRFs) to quantify perceptual sensitivity (d′) across all attention cueing and stimulation conditions. When the distractor was stimulated, exogenous attention yielded a characteristic response gain at the target—with performance benefits for valid cues and costs for invalid cues at high contrast levels. Crucially, this response gain was entirely preserved when the target was stimulated. This pattern contrasts with previous findings demonstrating that TMS to V1/V2 eliminates exogenous attentional effects at the stimulated location. These results indicate that rFEF+ is not necessary for exogenous attention, and together with previous studies [3-5] complete a double dissociation: rFEF+ is critical for endogenous but not exogenous attention, whereas V1/V2 is critical for exogenous but not endogenous attention. These findings reveal a distinct causal cortical architecture for voluntary and involuntary spatial attention, suggesting that they rely on different cortical scaffolds.

---

Given the high metabolic cost of cortical computation, the human brain must selectively process sensory information. Visual attention is a fundamental process enabling the brain to optimize its resources [1, 6, 7]. Via a push-pull mechanism [8], covert attention enhances signal processing at relevant locations at the expense of unattended locations—improving visual processing at the attended location (benefits) while suppressing signals at unattended locations (costs) [9-15]. There are two types of covert spatial attention: endogenous and exogenous. Endogenous attention is voluntary, goal-driven, sustained, and requires approximately 300 ms to be deployed. In contrast, exogenous attention is involuntary, stimulus-driven, and deployed rapidly and transiently (peaking at ∼100–120 ms) [for reviews, see [1, 2].

Both endogenous and exogenous attention distinctly improve performance across a wide array of visual tasks. Although their behavioral effects are often comparable, there are fundamental differences between them, particularly in how they modulate contrast sensitivity [13, 16-18] and spatial resolution [2, 19-22]. These differences manifest in tasks such as orientation discrimination and texture segmentation. For example, in a texture segmentation task, endogenous attention adjusts spatial resolution to improve performance [19, 23, 24], but exogenous attention reflexively enhances resolution even when doing so is detrimental to performance [e.g., 24-26]. Additional evidence shows that these two types of attention can also be behaviorally dissociated in other tasks [27-31].

Converging evidence from neurophysiological and neuroimaging studies indicates that covert spatial attention affects contrast sensitivity [32-37, for reviews, see 38-41]. Because orientation discriminability is contingent upon contrast sensitivity [42], performance is more accurate when covert attention is allocated to the stimulus location rather than elsewhere [1, 11-13, 43].

Functional magnetic resonance imaging (fMRI) studies of endogenous and exogenous attention have characterized partially overlapping networks in frontal and parietal lobes, often assuming similar effects in striate and extrastriate areas [44-47], but see [37]. Relevant to the current study, the right frontal eye field (rFEF+), the putative human homolog of the right macaque frontal eye field [48-50], is engaged during both endogenous [46, 47, 51-55] and exogenous [45, 56-59] attention. However, as a correlational technique, fMRI cannot establish whether a given region is functionally necessary, as it only records—but does not manipulate—neural activity.

TMS overcomes this limitation by manipulating neural activity—briefly disrupting excitation and inhibition—to enable causal inferences regarding the functional necessity of specific brain areas for perception and cognition [60, 61]. Critically, the behavioral impact of TMS depends on the initial activation state of the stimulated region. By manipulating brain states via psychophysical protocols—such as adaptation, covert attention, presaccadic attention, or priming—researchers can make informed predictions about how stimulation will impact behavioral performance [3-5, 62-72]. Specifically, TMS tends to suppress activity in neuronal populations that are currently active, while disinhibiting populations that are quiescent or less active.

Accordingly, TMS suppresses activity at attended locations and disinhibits activity at unattended locations. Leveraging this state dependency, prior TMS studies have established that early visual areas (V1/V2) are critical for exogenous attention, as stimulation eliminates its behavioral benefits and costs [3, 4], but are not critical for endogenous attention [5]. Conversely, TMS applied to the rFEF+ significantly diminishes the effects of endogenous attention [5]. Taken together, these findings raise an unresolved question central to establishing a formal double dissociation between the neural substrates of these two types of covert spatial attention: Does the rFEF+ play a critical role in exogenous attention? In the present study, we stimulated rFEF+ while manipulating the attentional state—using the same TMS-psychophysical protocol as in our previous studies [3-5]— to assess whether disrupting rFEF+ activity modulates the benefits and costs of exogenous attention.

## Results

### TMS Effects on Exogenous Attention

Participants performed an orientation discrimination task under six experimental conditions (3 attention x 2 stimulation) that were randomized across trials [**Figure 1**]. After a peripheral cue (valid, neutral, or invalid), two stimuli were presented simultaneously, one in the stimulated region, and the other in the symmetric region in the opposite hemifield. On every trial, participants received double-pulse TMS locked to the onset of the stimuli. Shortly after, participants reported the orientation of the stimulus indicated by a response cue, which either matched the peripheral cue (valid) or did not (invalid). This target coincided with the TMS-stimulated side (target–stimulated) or did not (distractor–stimulated). Importantly, stimulus onset was timed to coincide with the peak of exogenous spatial attention (∼80–120 ms after cue onset [1, 2, 21, 73]), and TMS was locked to stimulus presentation. The peripheral cue was uninformative regarding target location [3, 11, 74, 75], and participants were explicitly instructed of this fact.

**Figure 1.**
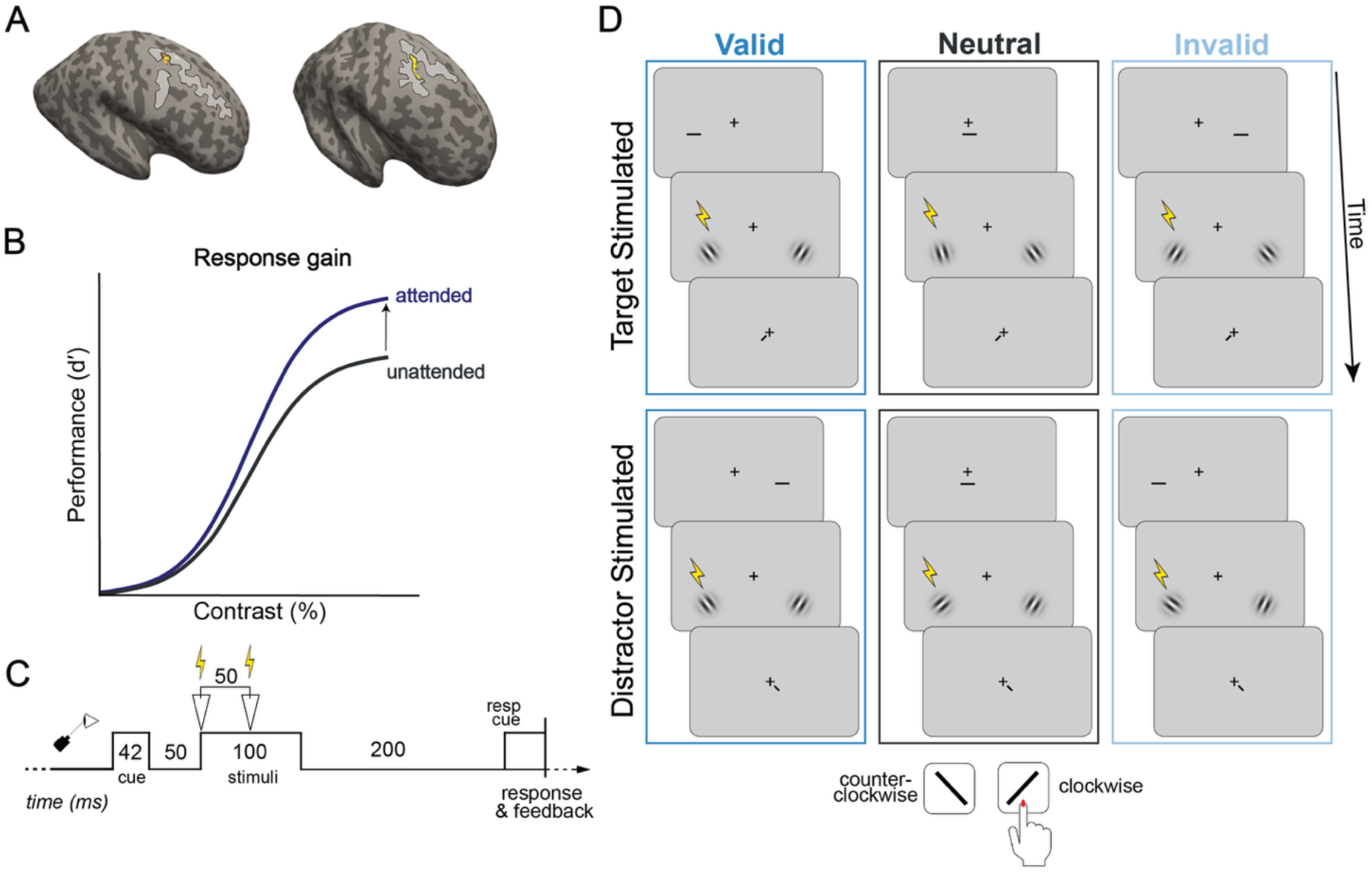
Psychophysics-TMS Task. **(A)** rFEF+ localization: Participants were stimulated over rFEF+ (yellow ROI), which was localized on each individual participant’s anatomy using the Wang et al. 2015 Atlas and validated via anatomical landmarks (the junction of the precentral and superior frontal sulcus (gray outlines). **(B)** Exogenous attention modulates contrast response functions via response gain: attention scales the response by a multiplicative gain factor—resulting in an increase in the asymptotic response (d′ _max_)–the maximal response achieved at high contrast). **(C)** Trial timeline: Participants received double-pulse TMS (50-ms inter-pulse interval) locked to stimulus onset. **(D)** Orientation discrimination task. In all trials, a central cue preceded stimulus presentation, indicating the target location (valid, 33% of trials), the distractor location (invalid, 33% of trials), or both stimulus locations (neutral 33% of trials). At the end of each trial, a response cue indicated the target Gabor whose orientation had to be reported; in valid trials, the location indicated by the cue and the response cue matched, whereas in invalid trials they did not match, and in neutral trials the response cue was equally likely to point to either location. In the target-stimulated condition, the response cue matched the stimulated region. In the distractor-stimulated condition, the response cue pointed to the location opposite the stimulated region. Participants indicated whether the stimulus was tilted clockwise or counterclockwise.

To estimate psychometric contrast-response functions (CRFs), which characterize the non-linear, sigmoidal relation between the contrast of a visual stimulus and the resulting neural or behavioral response [13, 14, 76, 77], we varied six stimulus contrast levels across trials. We then analyzed perceptual sensitivity (d′) for all 6 combinations of attention (valid, neutral or invalid) and TMS (target-stimulated, distractor-stimulated) conditions.

As in our previous studies [3-5], we expected the characteristic response gain associated with exogenous attention [12-14] in the distractor-stimulated condition: an increase in the upper asymptote following the valid cue (benefits) and a decrease following the invalid cue (costs), relative to the neutral cue. If rFEF+ is functionally necessary for exogenous attention, then disrupting its activity in the target-stimulated condition should significantly reduce this response gain by bringing the valid and invalid CRFs toward the neutral CRF.

### The effect of rFEF+ stimulation on Exogenous Attention

CRFs were obtained by fitting the performance data with Naka–Rushton functions [**Figure 2A**]. To analyze the causal role of the rFEF+ in exogenous attention, we conducted two Linear Mixed Models (LMMs), treating Attentional Cue (valid, neutral, invalid) and Stimulation Condition (target-stimulated, distractor-stimulated) as fixed effects, including their interaction, with “Participant” as a random intercept. For the upper asymptotic performance, d′_max_, there was a main effect of attentional cue [χ^2^(1)=13.22, p=.001; R_sp_^2^=.053]. Post-hoc comparisons revealed that d′_max_ was higher in the valid than in the neutral condition [t(11)=2.63, p=.024; Cohen’s *d*=0.76], which in turn was significantly higher than in the invalid condition [t(11)=4.27, p=.001; Cohen’s *d*=1.23]. There was no significant main effect of stimulation condition [χ^2^(1)=0.07, p=.786; R_sp_^2^=.002; BF_01_=8.18] nor a significant interaction [χ^2^(2)=0.52, p=.770; R_sp_^2^=.006; BF_01_=55.45]. For the semi-saturation constant, c_50_, neither attentional cue [χ^2^(2)=2.89, p=.236; R_sp_^2^=.029; BF_01_=17.01] nor stimulation condition [χ^2^(1)=0.42, p=.516; R_sp_^2^=.000; BF_01_=6.87] yielded significant main effects, nor did their interaction [χ^2^(2)=0.8, p=.669; R_sp_^2^=.007; BF_01_=48.19].

**Figure 2.**
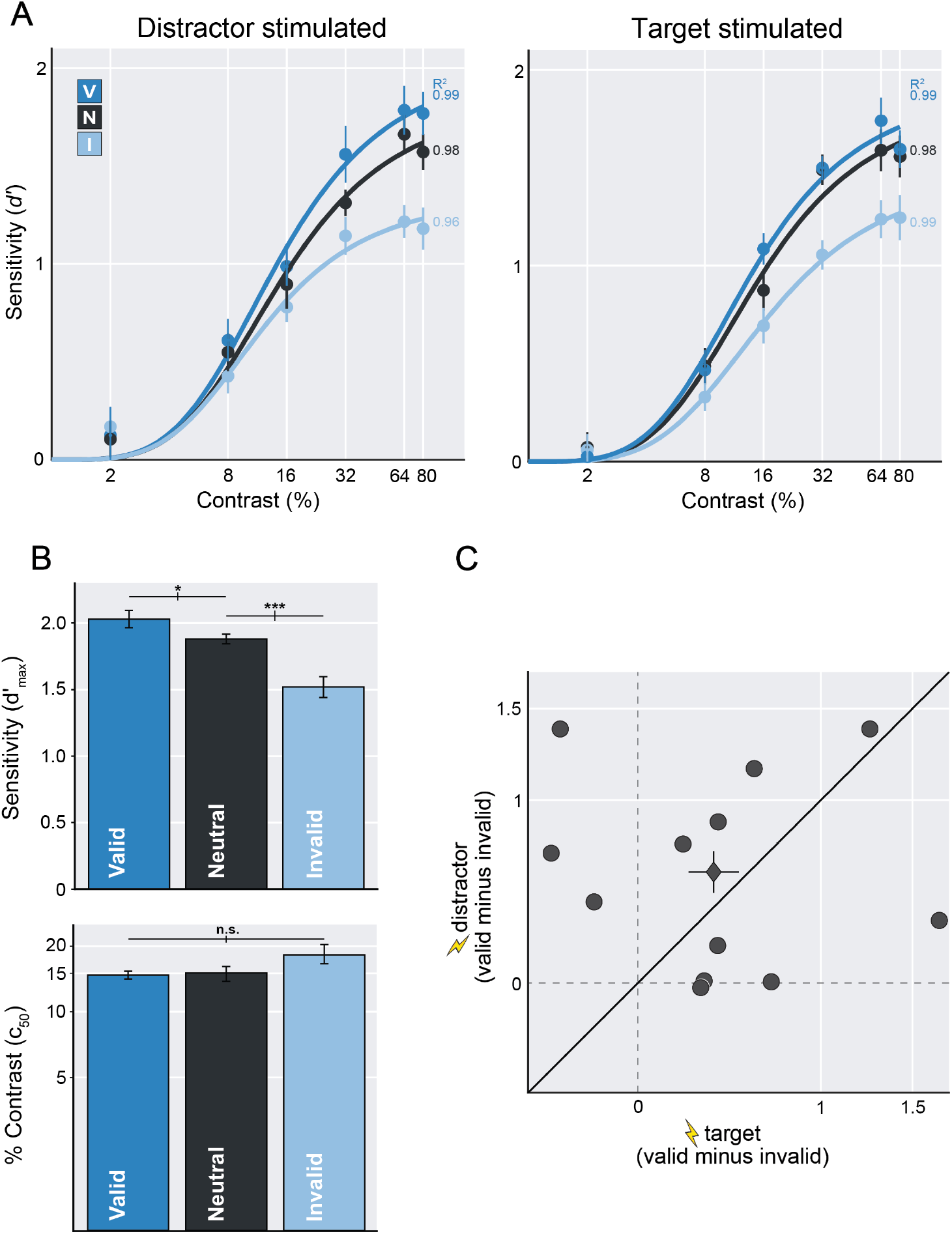
Contrast response functions for frontal (rFEF+) stimulation. **(A)** CRFs and parameter estimates for the upper asymptote d′_max_ and the semisaturation constant c_50_ in the distractor-stimulated condition (left panel) and the target-stimulated condition (right panel). **(B)** Mean parameter estimates for d′_max_ and c_50_ for their respective conditions. **(C)** Attentional effect (overall effect) computed as the difference in parameter estimates for d′_max_ between the valid and invalid conditions for the target-stimulated and distractor-stimulated conditions for each participant. Circles denote individual data; diamond denotes group mean. Error bars represent ±1 SEM. ***P≤0.001, *P≤0.05; Lightning bolts indicate the distractor-stimulated and the target-stimulated conditions.

Statistical effects are presented as mean parameter estimates for d′_max_ and c_50_ across both the distractor-stimulated and target-stimulated conditions **[Figure 2B]**. Individual data points for the overall attentional effect, calculated as the difference in d′_max_ estimates between valid and invalid conditions, are shown for each participant, comparing the target-stimulated versus distractor-stimulated trials **[Figure 2C**, see also **Figure S1B]**.

### Comparing rFEF+ and Occipital Stimulation for Exogenous Attention

In a previous study, using the same task and TMS protocol [3], we showed that occipital stimulation eliminates the effects of exogenous attention on performance. To directly compare the causal roles of rFEF+ and V1/V2 in exogenous attention, we performed a LMM analysis. We modeled the attentional effect (d′_max_ valid – d′_max_ invalid) as the dependent variable, with TMS Site (rFEF+,V1/V2), Stimulation Condition (target-stimulated, distractor-stimulated), and their interaction as fixed effects; a random intercept was included for each participant. The LMM revealed a significant interaction between TMS Site and Stimulation Condition (χ^2^(1)=3.86, p=.049; R_sp_^2^=.079). Simple effects analysis confirmed that the attentional effect was significantly larger in the distractor-stimulated than the target-stimulated condition when stimulating V1/V2 (p=.002; Cohen’s *d*=1.57), whereas no such difference emerged when stimulating rFEF+ (p=.328; BF_01_=3.43).

To further illustrate the impact of TMS Condition, individual data points for the overall attentional effect are shown for each participant, comparing the target-stimulated versus distractor-stimulated trials in which TMS was applied to V1/V2 in Fernández and Carrasco (2020) [3; **Figure S1A]**.

### Comparing rFEF+ Stimulation for Exogenous Attention and Endogenous Attention

In a previous study, using the same TMS protocol with the endogenous attention version of the same psychophysical task [5], we showed that rFEF+ significantly diminished the effects of endogenous attention on performance. To compare the causal role of the rFEF+ in both exogenous and endogenous attention, we performed a LMM analysis. We modeled the attentional effect (d′_max_ valid–d′_max_ invalid) as the dependent variable, with Attention Type (Exogenous, Endogenous), Stimulation Condition (target-stimulated, distractor-stimulated), and their interaction as fixed effects; ‘Participant’ was included as a random intercept. The LMM revealed a non-significant interaction between Attention Type and Stimulation Condition (χ^2^(1)=1.81, p=.178; R_sp_^2^=.024; BF_01_=2.80). However, planned comparisons revealed that the endogenous attention effect was significantly larger in the distractor-stimulated than the target-stimulated condition (p=.018; Cohen’s *d=*1.17), but no such difference was observed for exogenous attention (p=.326; Cohen’s *d=*0.41; BF_01_=3.43).

To further illustrate the effect of the attention type, individual data points for the overall attentional effect are shown for each participant comparing the target-stimulated versus distractor-stimulated trials in which TMS was applied to rFEF+ during endogenous attention in the study by Fernández, Hanning & Carrasco (2023) [5] **[Figure S1C]**.

### Microsaccade Frequency and Directionality

Given the well-established role of the rFEF+ in oculomotor control [49, 78] and our previous finding that TMS over the rFEF+ increases the rate of microsaccades directed toward the stimulated hemifield [5], we examined microsaccade frequency and directionality as a function of TMS stimulation. Microsaccade rates for both stimulation conditions showed the expected temporal dynamics across the trial sequence **[Figure 3A]:** a gradual decrease in anticipation of the cue that continued through cue presentation, with a dip at **∼**180 ms coinciding with stimulus presentation. This was followed by a sharp rise peaking at **∼**200 ms and then a decline within ∼100 ms. Approximately 100 ms after target onset (and the simultaneous TMS pulse), we observed a typical post-target rebound [5, 79-89]. Additionally, ∼100 ms after response cue presentation, we observed a response-cue rebound peaking at 600 ms [86, 88, 90]. A cluster-based permutation analysis (see *Methods: Microsaccade Analysis*) indicated that there were no significant differences in microsaccade rates between the two stimulation conditions.

**Figure 3.**
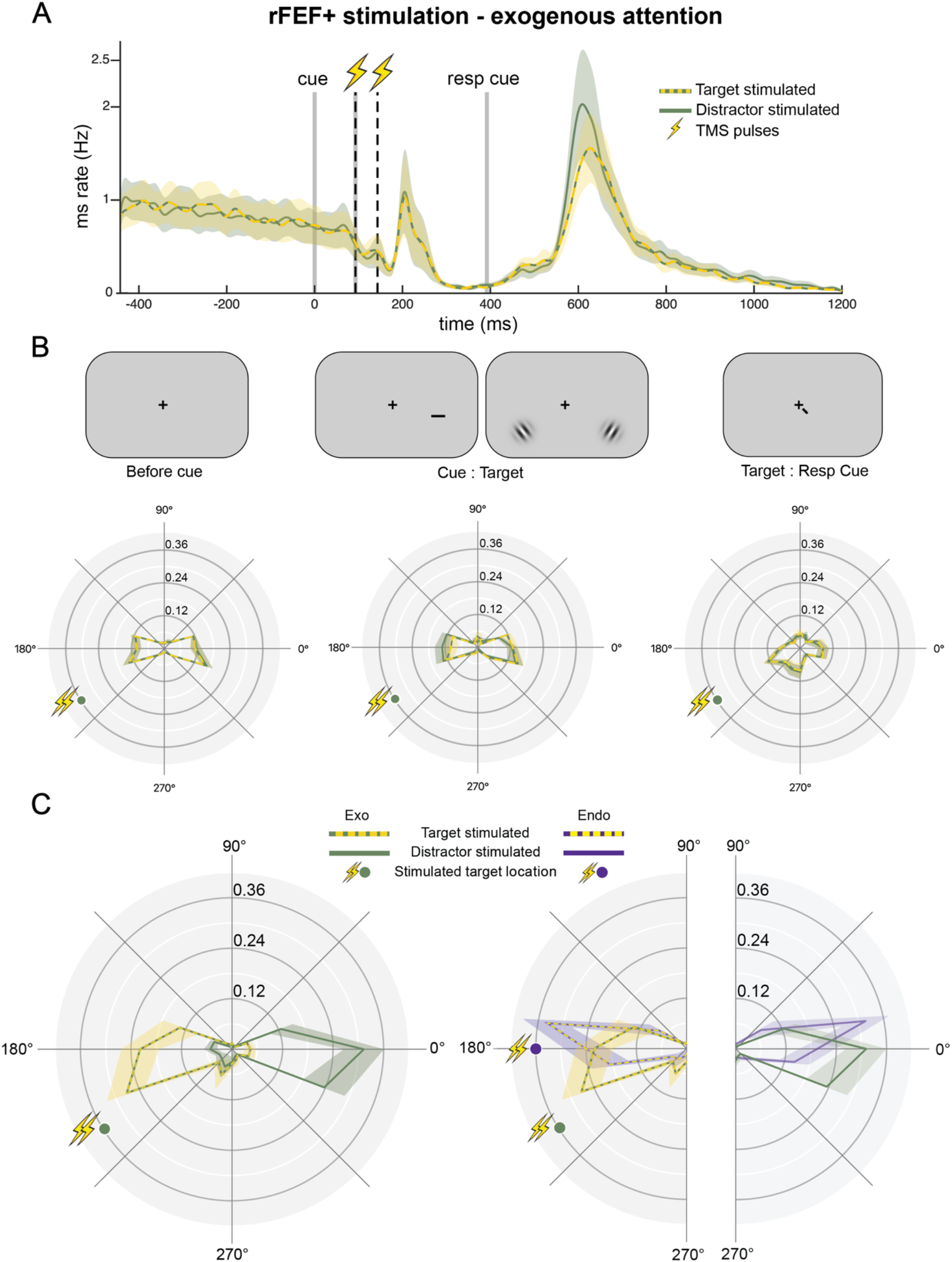
Microsaccade frequency and direction during frontal stimulation. **(A)** Group-averaged microsaccade rate relative to cue onset, split for target location (target stimulated, distractor stimulated). Vertical gray lines denote stimulus timing: cue onset, stimulus onset, and response cue onset, and vertical dashed black lines denote TMS pulses. **(B)** Normalized group-averaged polar angle microsaccade frequency before the cue (left panel), from cue onset to target onset (middle panel), and between target onset and response cue (right panel). **(C)** Normalized group-averaged polar angle microsaccade frequency after response cue onset (up to response) following rFEF+ TMS, split for target location (target-stimulated and distractor-stimulated conditions); current rFEF+ Exo data (left panel) and comparison with rFEF+Endo data reported in Fernández, Hanning & Carrasco (2023) [5] (right panel). Dots indicate target location in stimulated hemifield (green: Exo rFEF+; purple Endo rFEF+). Colored, shaded areas in all panels indicate ±1 SEM.

Regarding directionality [**Figure 3B**], microsaccades exhibited a largely symmetrical distribution in both the target-stimulated and distractor-stimulated conditions across all time windows preceding the response cue. We note that there were more microsaccades distributed toward the horizontal meridian than the vertical meridian. This pattern likely reflects the well-known horizontal bias in microsaccade direction [91, 92]. We evaluated the proportion of microsaccades directed toward the stimulated region (±35° polar angle relative to target location) following the onset of the response cue **[Figure 3C]** for 10 of the 12 participants (see Methods: *Microsaccade Analysis*).

We compared microsaccade directionality with our previous study, in which rFEF+ stimulation was combined with endogenous attention [5]; note that the target location differed [see Methods]. We used LMM to model microsaccade proportion as the dependent variable, with Attention Type (Exogenous, Endogenous), Stimulation Condition (target-stimulated, distractor-stimulated), and their interaction as fixed effects; “Participant” was included as a random intercept. The LMM revealed neither a significant interaction between Attention Type and Stimulation Condition (χ^2^(1)=0.32, p=.571; R_sp_^2^=.007; BF_01_=5.65) nor a main effect of Attention Type (χ^2^(1)=0.63, p=.426; R_sp_^2^=.013; BF_01_=4.83). However, there was a main effect of Stimulation Condition: the proportion of microsaccades directed toward the target was higher for the target-stimulated than the distractor-stimulated condition (χ^2^(1)=39.13, p<.001; R_sp_^2^=.568).

## Discussion

We manipulated exogenous attention using a well-established psychophysical protocol [9, 11-15, 18, 32, 93-98] and applied concurrent non-invasive brain stimulation (TMS) to probe its modulatory effects on visual perception [3, 4]. We investigated whether intact neural activity in rFEF+ is required for exogenous attention to modulate performance by assessing how TMS alters the contrast-response function (CRF). Consistent with previous psychophysical studies conducted without TMS [1, 11-14, 18, 32, 93], we observed significant benefits at the attended location and costs at the unattended location–via a response gain mechanism–when we stimulated the cortical representation of the distractor. This tradeoff in processing is consistent with a push-pull mechanism [8, 10-14]. Critically, when the target representation was stimulated, the magnitude of these benefits and costs remained entirely preserved.

Our novel findings reveal that although rFEF+ may be recruited during exogenous attention tasks– as demonstrated by fMRI studies [45, 56-58, 68]–this area is not functionally necessary for the modulatory effects of exogenous attention on contrast sensitivity. Rather than reflecting a contradiction, this discrepancy between fMRI and TMS results suggests that rFEF+ activation during exogenous orienting indexes processes that are not required for perceptual benefits. Consequently, our findings refine the functional map of the frontoparietal attention network, isolating rFEF+ as a site primarily dedicated to the top-down control of voluntary attention.

This finding stands in stark contrast to previous evidence demonstrating that both the benefits and costs of exogenous attention are eliminated when TMS is applied to V1/V2 [3, 4]. Because we used the same experimental protocol, task, and TMS pulse timing, we can rule out procedural factors as being responsible for the divergent effects observed when stimulating these two distinct cortical sites. Consequently, our results suggest that the early visual cortex—rather than rFEF+— serves as the critical functional substrate for the manifestation of exogenous attentional effects on perception. Although the frontoparietal network is involved in many aspects of spatial priority [5, 72, 99-108], its involvement may operate on a slower temporal scale. Critically, for endogenous attention, the roles of V1/V2 and rFEF+ are reversed: TMS applied to rFEF+ significantly diminishes the perceptual benefits of endogenous attention, whereas stimulation of V1/V2 does not modulate the magnitude of either benefits or costs [5].

Thus, with three previous findings—Exogenous and V1/V2 (modulation present) [3, 4]; Endogenous and V1/V2 (modulation absent) [5]; and Endogenous and rFEF+ (modulation present) [5]— the present findings (Exogenous and rFEF+; modulation absent) establish a formal double dissociation: V1/V2 is required for exogenous but not endogenous attention, whereas rFEF+ is required for endogenous but not exogenous attention. This double dissociation [**Figure 4**] provides compelling causal evidence that these two types of covert spatial attention rely on distinct, separable cortical scaffolds.

**Figure 4.**
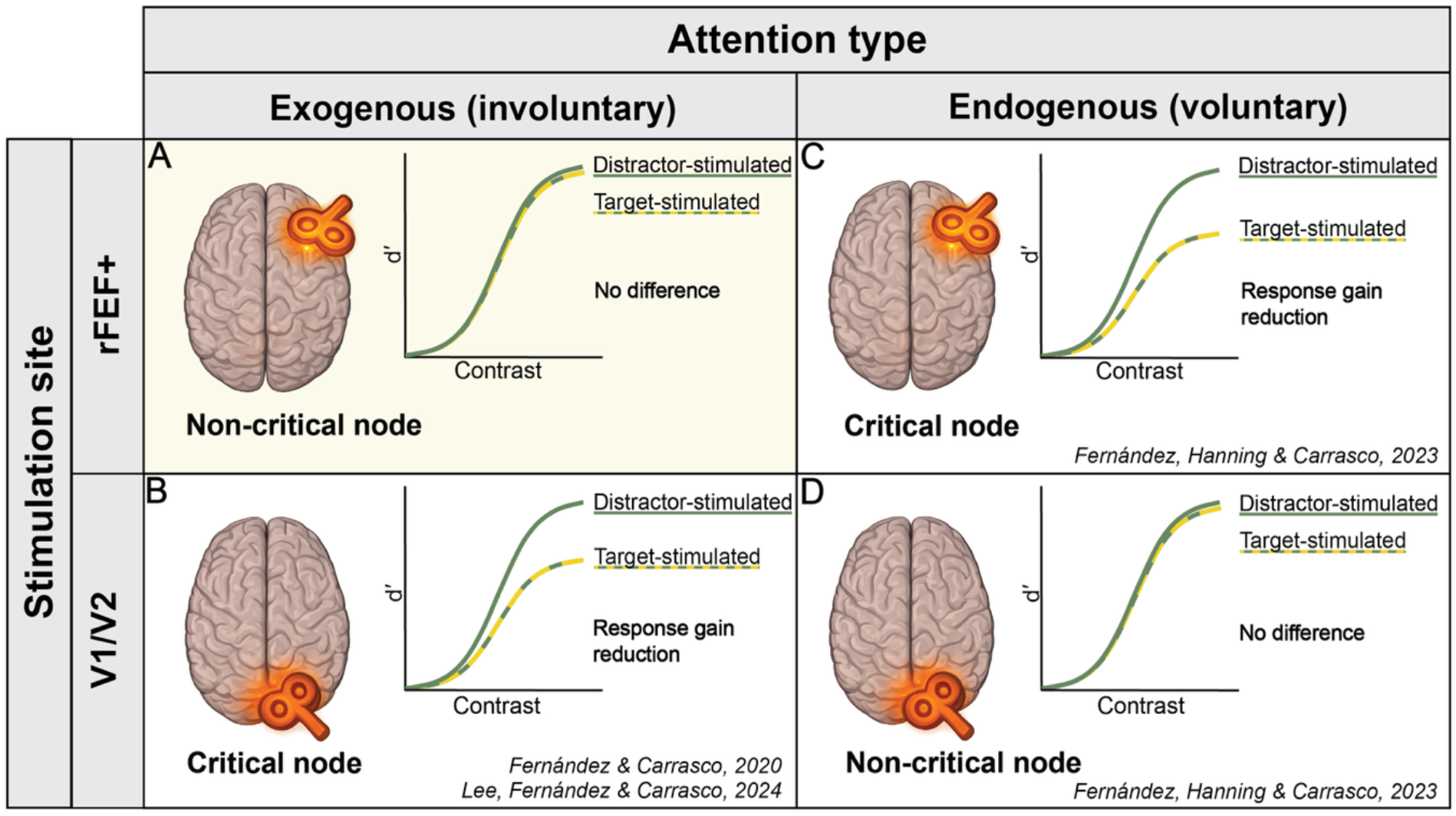
Double Dissociation Between Exogenous and Endogenous Attention. Summary matrix illustrating the causal necessity of distinct cortical regions in endogenous (voluntary) and exogenous (involuntary) covert spatial attention. Columns denote the targeted cortical nodes (frontal rFEF+ versus occipital V1/V2); rows denote attention type (Exogenous versus Endogenous). Within each quadrant, stylized brain maps indicate the targeted site of non-invasive TMS (orange icons); Contrast Response Functions (CRFs) depict the associated effect on behavior. **Left Column: Exogenous attention. (A)** rFEF+ stimulation has no effect on attentional modulation (current study). **(B)** V1/V2 stimulation eliminates the modulatory effects of exogenous attention [3, 4]. **Right Column: Endogenous attention. (C)** rFEF+ stimulation significantly attenuates attentional benefits and costs [5]; **(D)** V1/V2 stimulation has no effect [5]. Together, these results establish a double dissociation: V1/V2 is required for exogenous but not endogenous attention, whereas rFEF+ is required for endogenous but not exogenous attention.

The dissociation whereby TMS to V1/V2 eliminated the benefits and costs of exogenous attention [3, 4], but not those of endogenous attention [5], aligns with evidence that these different types of covert attention recruit early visual areas through distinct neural pathways. Exogenous attention is thought to operate via a rapid, stimulus-driven “feedforward” sweep that immediately boosts response gain in V1/V2 [109-111]—a process highly susceptible to the disruptive effects of TMS [112, 113]. In contrast, endogenous attention involves slower, recurrent feedback loops in which V1/V2 is “reactivated” at later time points [114, 115], receiving top-down feedback signals from frontal regions such as the FEF [55, 116-121]. This account is further supported by evidence of an attentional gradient across the visual hierarchy, where the magnitude of attentional modulation increases progressively from striate to higher extrastriate areas [37, 46, 52, 122, 123]. Moreover, the neural effect of endogenous attention is not always evident in V1, even when observed in the extrastriate cortex [37, 46]. Thus, early visual areas may play a secondary, less critical role in endogenous attention. Although V1–alongside the rFEF– can be involved in endogenous attention [124] and visual search tasks [125-127], it is not the critical functional substrate for endogenous attentional effects on perception. The absence of a TMS effect over early visual areas on endogenous attention suggests that top-down signals may be more robust to transient sensory disruption, or that the gain mechanisms implementing them are less susceptible to the parameters of our V1/V2 stimulation.

The dissociation in which TMS to rFEF+ modulated endogenous, but not exogenous, attention aligns with the account that the causal involvement of frontal nodes is state-dependent [71]. During endogenous attention, which relies on active top-down control, rFEF+ functions as a critical control hub and is therefore susceptible to disruption. Conversely, during exogenous orienting, rFEF+ is less susceptible to perturbation, consistent with the idea that it operates in a different functional state—potentially becoming functionally decoupled from the task-relevant circuitry or serving as a non-critical node for that specific process.

As with any neurostimulation or lesion study, demonstrating that a region is functionally necessary for a given perceptual or cognitive function does not imply that it is the sole source of that effect. Indeed, various regions play distinct roles in endogenous versus exogenous attention. For instance, TMS to the superior parietal lobule (SPL) selectively impairs the voluntary shifting of attention while leaving reflexive exogenous capture unaffected [128, 129]. Conversely, TMS disruption of the right intraparietal sulcus (r-IPS) or the right temporoparietal junction (r-TPJ) abolishes exogenous attention benefits while leaving endogenous effects unaltered [130]. These findings, alongside our own, indicate that although both types of attention rely on a distributed network, they depend on distinct causal nodes within that network.

The microsaccade analysis revealed temporal dynamics consistent with previous reports: a gradual decrease in microsaccade rate in anticipation of the cue, which continued through cue presentation, followed by a sharp rise and a subsequent decline until response-cue onset. Shortly after target presentation (and the simultaneous TMS pulse), we observed a typical post-target rebound [5, 79-89]. A similar rebound followed response-cue presentation [86, 88, 90], likely reflecting the transition from stimulus-driven attentional engagement to the task-related requirements of the response phase. Critically, the directional analysis following the response cue closely resembled that observed during rFEF+ stimulation in endogenous attention [5]. In both cases, microsaccades were more frequently directed toward the target location following the onset of the response cue, consistent with a shift of attention toward the task-relevant stimulus. This directional effect —which may reflect participants’ attempts to retrieve the target information indicated by the response cue while preparing the response [131] —is consistent with findings in perception [88], attention [90], and visual short-term working memory [132].

Our findings challenge the premise that all shifts of spatial attention—even covert—depend on a common oculomotor substrate, a core tenet of the Premotor Theory of Attention [133-135]. This framework suggests that spatial attention is intrinsically inseparable from the neural systems responsible for motor preparation. However, our results reveal that rFEF+—a core node within the oculomotor control network—is not functionally necessary for exogenous attention, despite its established role in endogenous attention. This dissociation indicates that not all forms of covert attention depend on a shared oculomotor substrate. This conclusion is reinforced by previous research establishing that V1/V2 and rFEF+ play dissociable roles in covert endogenous attention [5] and exhibit a different pattern of dissociation during presaccadic attention [72]. Specifically, whereas rFEF+ stimulation enhances sensitivity opposite the saccade target throughout saccade preparation, V1/V2 stimulation reduces sensitivity at the saccade target only shortly before saccade onset. Furthermore, V1/V2 TMS at a fixed time (∼150 ms) after cue onset does not affect presaccadic attention—unlike covert exogenous attention, which is extinguished using the same psychophysical protocol and TMS pulse timing [3, 4]. Together, these findings show that occipital regions are recruited primarily at later stages of saccade preparation. Overall, the converging evidence indicates that exogenous, endogenous, and presaccadic attention rely on distinct patterns of involvement of V1/V2 and rFEF+ and, therefore, do not share a single, unified neural implementation as proposed by the Premotor Theory.

The current findings add to the mounting evidence that covert spatial attention and presaccadic attention are dissociable processes [136-138]; review, [139]. If exogenous attention were merely a byproduct of premotor activation in the frontal oculomotor system, disruption of rFEF+ should have abolished, or at least significantly attenuated, the exogenous attentional effect. Instead, our results demonstrate that exogenous attention can operate independently of rFEF+ activity. This finding is consistent with the view that stimulus-driven, involuntary covert shifts of attention may be phylogenetically more primitive, facilitating automatic responses to environmental demands and rapid reactions to potentially behaviorally relevant stimuli [1]. Whereas reflexive orienting is observed widely across the animal kingdom, sustained endogenous attention is primarily observed in cognitively flexible species [140]. These reflexive shifts likely rely on subcortical and posterior circuits, whereas endogenous orienting is governed by more recently evolved frontoparietal cortical networks [141].

In conclusion, our results reveal that rFEF+ is not a functionally necessary component of the neural circuitry underlying exogenous spatial attention. By establishing that rFEF+ stimulation does not modulate the costs and benefits of stimulus-driven attention—despite its demonstrated causal role in endogenous and presaccadic attention—we provide the final evidence for a double dissociation between frontal and occipital nodes of the covert spatial attention network. These findings indicate that exogenous attention can operate via a response gain mechanism independently of frontal involvement. Ultimately, this study refutes the necessity of a unified premotor substrate for all forms of spatial selection, suggesting instead that the human brain utilizes distinct, specialized cortical scaffolds to prioritize information depending on whether attention is guided by internal goals or captured by the environment.

## Methods and Materials

### Participants

Twelve participants (7 females; age range: 22-31 years, mean age=27.82, SD=3.29) with normal or corrected-to-normal vision participated in this experiment. All participants, except for two authors (Q.C. and H.H.L), were naïve to the study’s purpose. All participants were right-handed, provided written informed consent, and were screened for TMS contraindications. The experimental protocol adhered to TMS research safety guidelines and was approved by the Institutional Review Board for Human Subjects at New York University. All participants had their corresponding T1-weighted anatomical images archived in the NYU neuroimaging database.

### Apparatus

All participants completed the experiment in a dark room, with their heads stabilized by a chin-and-forehead rest at a viewing distance of 55 cm from a gamma-calibrated ViewPixx/EEG LCD monitor (1,920 x 1,080 resolution, 120 Hz refresh rate; calibrated with a ColorCAL MKII colorimeter, Cambridge Research Systems). Stimuli were generated and displayed using MATLAB (MathWorks) and the Psychophysics Toolbox [142-144]. A Linux desktop machine controlled both stimulus presentation and the collection of behavioral responses.

Participants viewed the stimuli binocularly. To ensure the manipulation of covert spatial attention, gaze position of their dominant eye was monitored at a sampling rate of 1,000 Hz using a desktop-mounted SR Research Eyelink 1000 eye tracker (SR Research, Osgoode, ON, Canada). Due to a technical error, eye-tracking data from two participants were lost and could not be recovered.

Experimental trials were gaze-contingent to ensure stable fixation: any trial in which gaze deviated more than 1.5° of visual angle from fixation, or during which a blink was detected, was immediately aborted and re-queued to the end of the experimental block.

### Stimuli

A black fixation cross (subtending 0.7° x 0.7°of visual angle, composed of two perpendicular lines) was displayed in the center of the screen throughout the experiment on a uniform gray background. The stimuli in each trial consisted of two Gabor patches (2-cpd sinusoidal gratings embedded in a Gaussian envelope). The Gabor stimuli subtended 3.83° of visual angle. They were centered at 9.43° of eccentricity (5° below fixation and ± 8° horizontally from fixation), approximating the location in our previous exogenous and endogenous attention TMS studies [3, 4]. Stimulus contrast was defined as Michelson contrast and sampled from six levels (2%, 8%, 16%, 32%, 64%, and 80%). The peripheral attention cue was a black horizontal line (2° in length, 0.12° in thickness), positioned 1° below and horizontally centered on the cued Gabor, whereas the neutral cue was 1° below fixation. The response cue consisted of a black line (0.58° in length, 0.14° in thickness) oriented in the direction of the target (lower left or lower right).

### TMS Machine and Neuronavigation

Participants were stimulated with an MCF-B70 coil positioned over the frontal cortex, with the coil handle oriented perpendicular to the sagittal plane. TMS pulses were generated using a MagPro X100 stimulator (MagVenture, Farum, Denmark) and triggered via a MATLAB–Arduino interface. For each participant, the stimulation intensity was calibrated to the individual resting motor threshold and remained constant across the experimental session. Across all participants, stimulation intensity ranged from 51% to 59% (mean=54.37%, SD=2.74%) of the maximum stimulator output. Coil placement was guided using the neuronavigation software Brainsight (Rogue Research, Montréal, QC, Canada) by importing each participant’s anatomical T1-weighted MRI image with a predefined region of interest (ROI) corresponding to the right frontal eye field (rFEF+). Brainsight was used to monitor and record coil position on the scalp with millimeter-level precision, ensuring consistent targeting of the same cortical site across experimental sessions.

### Titration procedure

To equate task difficulty across participants and to minimize learning effects, we independently titrated the tilt angles for leftward and rightward Gabor stimuli for each participant at the beginning of every psychophysics–TMS session (without TMS) using an adaptive staircase procedure [145] implemented in the Palamedes toolbox [146]. We used a neutral condition with high contrast (80%) stimuli to determine the tilt angle at which each participant’s orientation discrimination performance reached a 75% accuracy threshold. This individualized tilt angle was then used in the psychophysics–TMS session across all conditions (left Gabor group average: 0.89° ± 0.09°, right Gabor group average:0.94° ± 0.10°; mean±SEM).

### Psychophysics-TMS Task

Participants completed five psychophysics–TMS sessions, each lasting approximately 1.5 hours. At the beginning of every session, an adaptive staircase procedure (see *Titration procedure*) was used to control for potential learning effects. During each session, participants performed a two-alternative forced-choice (2AFC) orientation discrimination task **[Figure 1C,D]**. Each trial began with a variable fixation window (750, 1,150, or 1,750 ms), followed by a peripheral attention cue (valid, neutral, or invalid) presented for 41.67 ms. All cues were equally likely and therefore uninformative regarding target location and orientation. After a 50 ms blank period, two Gabor patches were presented for 100 ms. We employed a double-pulse TMS protocol (see *TMS protocol*), where the first pulse was time-locked to Gabor onset and the second followed 50 ms after. The interval between cue onset and stimulus presentation was chosen to maximize the deployment of exogenous attention. After another 200 ms blank interval, a response cue was presented, indicating the target stimulus.

If the response-cued (target) location matched the cue, the trial was considered valid; if it did not match, it was considered invalid. Importantly, if the response-cued target was in the stimulated region the trial was labeled “target–stimulated,” otherwise it was labeled as “distractor–stimulated.” The response cue remained on the screen until a response was recorded. Auditory feedback (400 Hz., 150 ms, normalized amplitude 0.2) was provided for incorrect trials. Each participant completed a total of 3,600 trials: 100 trials × 6 contrast levels × 3 Attentional Cues (valid, neutral, and invalid) × 2 Stimulation Conditions (target and distractor).

This design ensured an identical stimulation protocol across all experimental conditions and thus provides a stronger control than alternative protocols, such as sham stimulation, which only reproduce some of the sensations produced by TMS [147, 148].

### TMS protocol

A double-pulse TMS was administered on each trial, with a 50 ms inter-pulse interval. This stimulation protocol was chosen to optimize direct comparisons with our previous exogenous attention manipulation using TMS over V1/V2 [3], as well as with our previous endogenous attention manipulation using TMS over V1/V2 or to rFEF+ [5]. It is also widely adopted in studies investigating visual perception and attention [124, 125, 127, 130], as well as motor functions [149-151], to probe intracortical circuitry, including both inhibitory (GABAergic) and excitatory (glutamatergic) pathways. Moreover, this timing is consistent with a long-interval intracortical inhibition (LICI) protocol, a method established to produce robust interference with ongoing neuronal processing [152, 153].

### Anatomical Data Acquisition

For each participant, raw full-brain anatomical data were obtained from the NYU Retinotopy Database [154]. Anatomical scans were collected at the NYU Center for Brain Imaging on a 3T Siemens MAGNETOM Prisma MRI scanner (Siemens Medical Solutions) using a Siemens 64-channel head coil. Each participant contributed one full-brain, T1-weighted (T1w) MPRAGE anatomical image (TR 2,400 ms; TE 2.4 ms; voxel size 0.8 mm^3^ isotropic; flip angle 8°). All T1w scans were aligned to a common template to ensure consistent slice prescriptions across participants, and cortical surfaces were reconstructed using the *FreeSurfer’s recon-all* pipeline [155].

### Frontal Eye Field Localization

The putative human right frontal eye field (rFEF+) was localized using the Wang atlas [156], and validated via anatomical landmarks (lying within the junction of the precentral and superior frontal sulci), as in previous studies [5, 72, 157]. The rFEF+ region of interest (ROI) was projected into each participant’s native volume using the *FreeSurfer*’s *mri_surf2vol* and *mri_surf2surf* functions [158]. Localization was further validated using anatomical landmarks at the junction of the precentral and superior frontal sulci [48, 50]. Each participant’s T1 image and rFEF+ ROI were then imported into Brainsight, enabling precise neuronavigation and accurate stimulation of rFEF+ across sessions.

### Electric field simulation

We conducted electric field (E-field) simulations using the simNIBS toolbox [159, 160] to validate that the selected stimulation parameters generated sufficient field strength at the intended target, confirming effective stimulation of rFEF+ [**Supplemental Figure 2**]. This approach also enabled direct comparison with previous studies applying TMS to the occipital cortex and rFEF+ [3-5], which used a different stimulator system (Magstim Rapid Plus stimulator (3.5T), with maximal current strength of 1.53 V/m). We confirmed that the stimulation over rFEF+ in the current study (maximal current strength of 1.96 V/m) was sufficient, even stronger than that in previous work.

### Task performance

Task performance was quantified by sensitivity, indexed by d’ [d’ = z(hit rate) − z(false-alarm rate)] and was measured as a function of stimulus contrast. Hits were defined as trials in which participants correctly discriminated the clockwise-tilted stimuli, whereas incorrect discriminations of counter-clockwise stimuli were classified as false alarms [3, 5, 14, 17, 98, 161-163]). We applied the log-linear correction method to avoid undefined values in cases of extreme hit or false alarms rates [164]. Performance was measured using the method of constant stimuli with six interleaved contrast levels. The contrast-response functions (CRF) for each participant was then modeled by fitting the data with a Naka–Rushton function [76]:

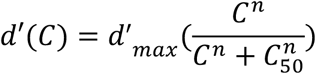

where d′(C) denotes sensitivity as a function of contrast, d′_max_ is the asymptotic performance level, c_50_ is the semisaturation constant (the contrast at which half-maximum performance is reached), and *n* determines the slope of the function. Fits were obtained by minimizing the residual sum of squares using a non-linear least-squares criterion. During optimization, d′_max_ and c_50_ were treated as free parameters, whereas *n* was fixed to the value that best fit the group-averaged data across all conditions. Contrast levels were log-transformed before fitting.

### Quantification and Statistical Analysis

For the analysis of CRF, we used a non-linear least-squares approach to fit a Naka-Rushton equation to the data, comparing the d′_max_ (response gain) and c_50_ (contrast gain) parameters across conditions. All reported t-tests are two-tailed, and error bars in figures represent within-participant SEM. Effect sizes for fixed effects and interactions in the LMM were estimated using semi-partial R2 (Rsp2) as implemented in the r2glmm package [165, 166]. Values of 0.02, 0.13 and 0.26 have been proposed as useful estimates of small, medium and large effect sizes [167].

To complement reported null effects, we conducted Bayesian model comparisons using the Bayesian Information Criterion (BIC) to derive Bayes factors [168]. Estimates of Bayes factors (BF_01_) were derived from BIC differences between nested LMM models fitted using maximum likelihood. Values of BF_01_ exceeding 3, 7, and 10 are considered positive, strong, and very strong evidence, respectively, in favor of the null hypothesis [169].

All Linear Mixed Models (LMMs) were fitted using the ‘lme4’ package in R (v4.3.0), with significance assessed via Satterthwaite’s approximation for degrees of freedom (‘lmerTest’ package). Significant interactions were followed by simple effects analyses using the ‘emmeans’ package, with p-values adjusted for multiple comparisons using the Holm-Bonferroni method. ‘Participant’ was included as a random intercept in all models to account for idiosyncratic differences in baseline sensitivity.

To assess the causal role of rFEF+ in exogenous attention, we fit two LMMs —one for d′max and one for c50— with Attentional Cue (Valid, Neutral, Invalid) and Stimulation Condition (target-stimulated, distractor-stimulated) as fixed effects, including their interaction.

To compare the causal roles of rFEF+ and V1/V2 in exogenous attention, we fit a LMM with attentional effect as the dependent variable, calculated as the difference in asymptotic performance between valid and invalid trials, (d′_max_ valid–d′_max_ invalid). We treated TMS Site (rFEF+, V1/V2), Stimulation Condition (target-stimulated, distractor-stimulated), and their interaction as fixed effects.

To compare the causal roles of rFEF+ in exogenous versus endogenous attention, we fit an LMM with attentional effect (d′_max_ valid–d′_max_ invalid) as the dependent variable. We treated Attention Type (Endogenous, Exogenous), Stimulation Condition (target-stimulated, distractor-stimulated), and their interaction as fixed effects.

### Microsaccade Analysis

Eye-movement data were collected at 1,000 Hz and analyzed offline in MATLAB (MathWorks). Microsaccades were identified using a velocity-based detection algorithm [170] and defined as saccades with an amplitude <1.5°. Onsets and offsets were determined when velocity exceeded or dropped below the median moving average by at least 6 SDs for a minimum duration of 6 ms. Only events separated by at least 10 ms were accepted as distinct microsaccades; detection was terminated at the onset of blinks or once the trial response was recorded. Microsaccade amplitude and peak velocity followed the expected “main sequence” relation and were comparable across experimental sessions.

For all included events (6,651 ± 1,175), we calculated microsaccade rate (Hz) within the interval from −750 to +1,500 ms relative to cue onset. Microsaccade onsets were averaged across trials at each sampling point for each condition and multiplied by the sampling rate to yield a rate time series. The resulting microsaccade–rate functions were subsequently smoothed with a 10-ms Gaussian kernel. To further assess the impact of TMS on microsaccade frequency, we employed a cluster-based permutation testing [171], which identifies temporal intervals of significant differences between two conditions while controlling for multiple comparisons. Consecutive time samples with p < 0.05 were combined into temporal clusters, and the largest cluster (defined by its summed *t-statistic* mass) was evaluated against a null distribution generated from 1,000 random permutations of condition labels within or across participants.

To assess microsaccade directionality, we categorized microsaccades according to their polar angle–defined between onset and offset positions–into 16 bins (22.5° per bin) centered on the cardinal axes and computed separately for each experimental condition. For each participant, the proportion of microsaccades in each bin was normalized by the total number of microsaccades recorded within that condition.

## ACKNOWLEDMENTS

This research was supported by NIH R01-EY019693 to MC, The Ministry of Education in Taiwan to HHL, Marie Skłodowska-Curie individual fellowship (MSCA-IF 898520) by the European Commission to NMH, and P30 EY013079 (Core Grant for Vision Research) to New York University. We thank past and present Carrasco Lab members for feedback, especially Laura Dugué and Antonio Fernández (who also collected preliminary data), and Klara Hoxha for assistance with data collection.

## AUTHOR CONTRIBUTIONS

MC conceived, funded and supervised the study; QC and HHL collected and QC, HHL and NH analyzed the experimental data; all authors interpreted the results; NH co-supervised the study; QC wrote the Methods; MC wrote the manuscript. All authors edited the manuscript and approved the final version.

## Supplemental Figures

**Figure S1.**
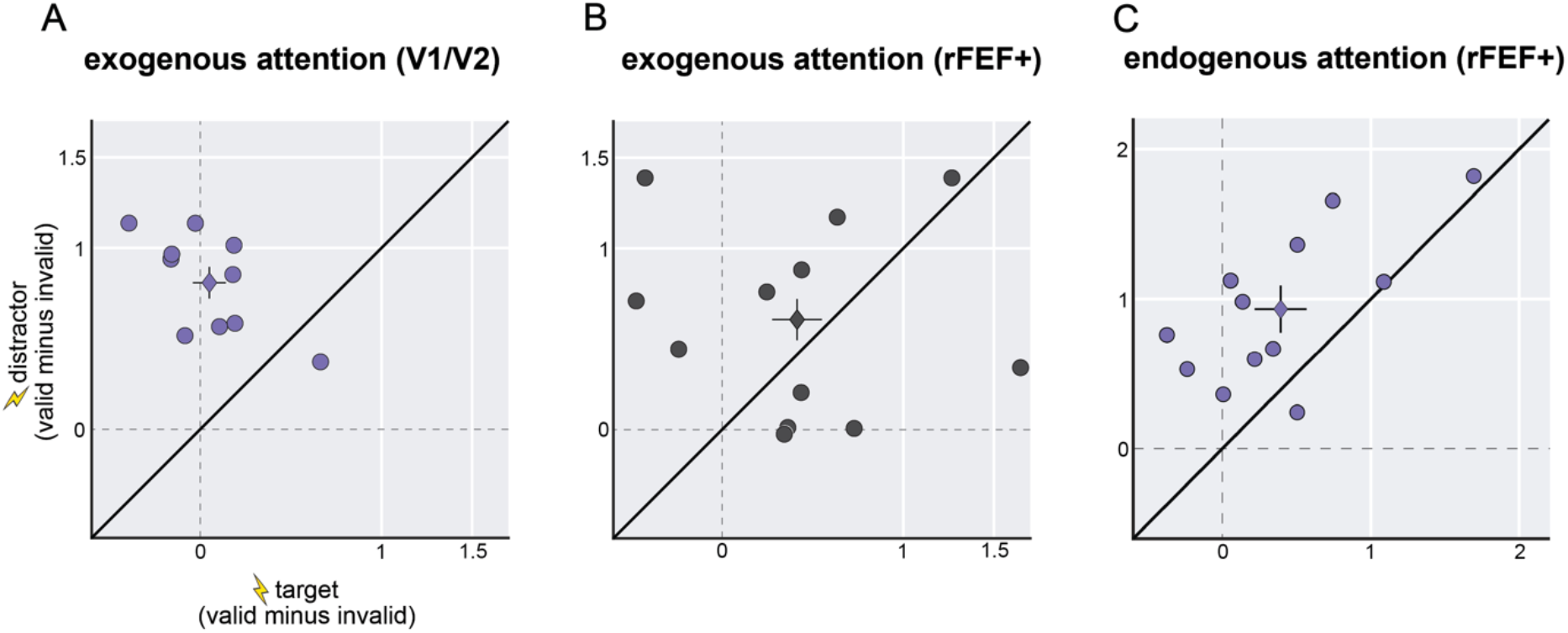
Individual attentional effects across studies. The overall attentional effect was calculated as the difference in d′_max_ parameter estimates between valid and invalid conditions for both the target-stimulated and distractor-stimulated conditions for each participant. (**A**) Data from Fernández and Carrasco (2020) involving exogenous attention and TMS over V1/V2 [3]. (**B**) Data from the current study involving exogenous attention and TMS over rFEF+. (**C**) Data from Fernández, Hanning and Carrasco (2023) involving endogenous attention and TMS over rFEF+ [5]. Circles represent individual data; diamonds represent the group mean.

**Figure S2.**
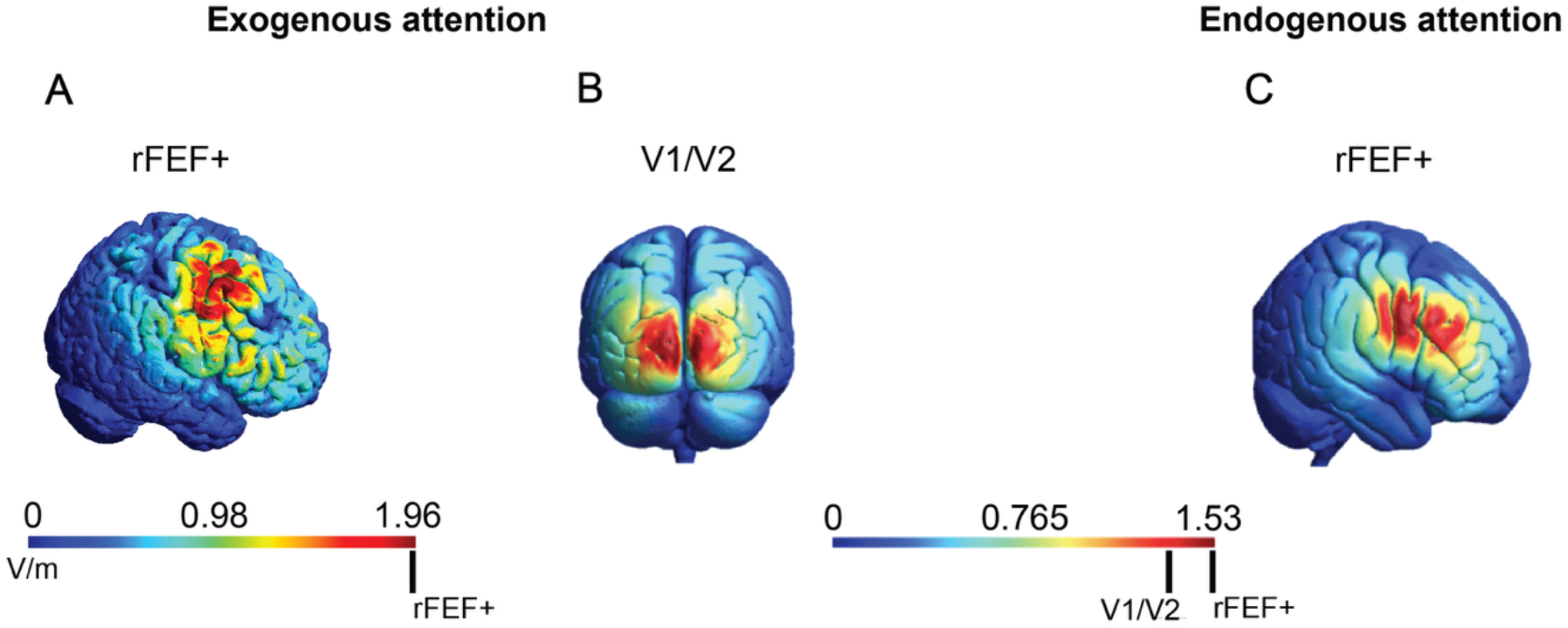
Simulated cortical excitability. Estimated E-field for the rFEF+ and occipital pole (V1/V2) using the simNIBS toolbox. **(A)** The E-field in the current study involving exogenous attention and TMS over rFEF+ is approximately 1.96 V/m. **(B)** The E-field in the Fernández and Carrasco (2020) involving exogenous attention and TMS over V1/V2 in Fernández & Carrasco (2020) [3] and in Lee, Fernández & Carrasco (2024) [4] is approximately 1.36 V/m. **(C)** The E-field in the Fernández, Hanning and Carrasco (2023) involving endogenous attention and TMS over rFEF+ [5] is approximately 1.53 V/m.

